# Biochemical characterization of naturally occurring mutations in SARS-CoV-2 RNA-dependent RNA polymerase

**DOI:** 10.1101/2024.02.24.581855

**Authors:** Matěj Danda, Anna Klimešová, Klára Kušková, Alžběta Dostálková, Aneta Pagáčová, Jan Prchal, Marina Kapisheva, Tomáš Ruml, Michaela Rumlová

## Abstract

Since the emergence of SARS-CoV-2, mutations in all subunits of the RNA-dependent RNA polymerase (RdRp) of the virus have been repeatedly reported. Although RdRp represents a primary target for antiviral drugs, experimental studies exploring the phenotypic effect of these mutations have been limited. This study focuses on the phenotypic effects of substitutions in the three RdRp subunits: nsp7, nsp8, and nsp12, selected based on their occurrence rate and potential impact. We employed nano-differential scanning fluorimetry and microscale thermophoresis to examine the impact of these mutations on protein stability and RdRp complex assembly. We observed diverse impacts; notably, a single mutation in nsp8 significantly increased its stability as evidenced by a 13 °C increase in melting temperature, whereas certain mutations in nsp7 and nsp8 reduced their binding affinity to nsp12 during RdRp complex formation. Using a fluorometric enzymatic assay, we assessed the overall effect on RNA polymerase activity. We found that most of the examined mutations altered the polymerase activity, often as a direct result of changes in stability or affinity to the other components of the RdRp complex. Intriguingly, a combination of nsp8 A21V and nsp12 P323L mutations resulted in a 50% increase in polymerase activity. Additionally, some of the examined substitutions in the RdRp subunits notably influenced the sensitivity of RdRp to Remdesivir®, highlighting their potential implications for therapeutic strategies. To our knowledge, this is the first biochemical study to demonstrate the impact of amino acid mutations across all components constituting the RdRp complex in emerging SARS-CoV-2 subvariants.

**Significance statement:** While the impact of SARS-CoV-2 spike protein mutations has been extensively explored, our understanding of mutations within the RNA-dependent RNA polymerase (RdRp), crucial for viral replication and a key target for antivirals like Remdesivir, remains limited with studies conducted solely *in silico*. We focused on selected RdRp mutations identified from December 2019 to June 2022, assessing their effects on enzyme stability, complex assembly, and activity. Advanced biochemical analyses reveal how these mutations can alter RdRp functionality, providing insights into viral evolution and resistance mechanisms. This study, pioneering in assessing the biochemical implications of RdRp mutations, provides invaluable insights into their roles in viral replication and antiviral resistance, hereby opening new pathways for developing therapies against the continuously evolving SARS-CoV-2 variants.

## Introduction

The Coronavirus Disease 2019 (COVID-19) emerged in December 2019 and shortly after escalated into a global pandemic. As of December 24, 2023, there have been 773,119,173 reported cases with 6,990,067 fatalities^1^. Although the World Health Organization has declared that COVID-19 is no longer a public health emergency of international concern, it continues to present a severe health threat, particularly to vulnerable groups^2^. The causative agent of COVID-19 is Severe Acute Respiratory Syndrome Coronavirus 2 (SARS-CoV-2). Typically, coronaviruses cause mild to moderate respiratory illnesses. However, specific strains within the *Betacoronavirus* genus have been associated with severe outbreaks, namely SARS-CoV, which led to the Severe Acute Respiratory Syndrome (SARS) outbreak in 2003, and MERS-CoV, known for the Middle East Respiratory Syndrome (MERS)^3^.

Betacoronaviruses are enveloped viruses with a positive-sense single-stranded RNA genome of approximately 30 kilobases (kb). This categorizes them among the largest known RNA viruses^4^. Their genome encodes four structural proteins, and multiple nonstructural and accessory proteins. A major part of the genome is represented by ORF1a and ORF1ab, which are translated into two large polyproteins pp1a and pp1ab. Upon translation, the pp1a and pp1ab polyproteins undergo cleavage, mediated by a papain-like protease, resulting in the production of 10 and 16 nonstructural proteins (nsps), respectively. The main role of these nonstructural proteins is to mediate the transcription and replication of viral RNA through various mechanisms. Out of all the 16 nsps, nine (nsp7 – nsp10; nsp12 – nsp16) assemble to form the coronaviral replication-transcription complex (RTC), which is directly responsible for RNA replication^5–7^.

Amidst other replication factors in the RTC, nsp12, a canonical RNA-dependent RNA polymerase (RdRp), possesses pivotal enzymatic activity. Structurally, nsp12 adopts a typical ‘right-hand’ architecture with subdomains corresponding to the thumb, palm, and fingers. These motifs, conserved amongst the RNA viruses^8^, collectively form the catalytic site of the polymerase. Furthermore, nsp12 includes a Nidovirus RdRp-Associated Nucleotidyl Transferase (NiRAN) domain participating in the capping process^9, 10^. On its own, nsp12 exhibits only minimal RdRp activity. It gains most of its catalytic activity upon binding to accessory subunits nsp7 and nsp8, forming minimal complex capable of mediating RNA replication, nsp7:nsp8:nsp12^11^. This RdRp complex contains a single nsp7 and two nsp8 subunits; nsp8a directly interacts with nsp12, while nsp8b is bound via nsp7. While nsp7 is a small protein consisting of three *α*-helices, each nsp8 subunit has a “golf club”-like shape, with a C-terminal head domain and a rod-like N-terminal extension that stabilizes both the newly synthesized RNA and template duplex (Fig. 1, A)^11, 12^. The RdRp of SARS-CoV-2, like other RNA viruses, operates with low intrinsic fidelity^13^. To replicate its 30 kb long genomic RNA effectively and avoid lethal mutagenesis, coronaviruses developed an elaborate mechanism utilizing a viral 3′–5′ exoribonuclease (nsp14) as an independent proofreading factor^4, 14^.

**Figure 1.**
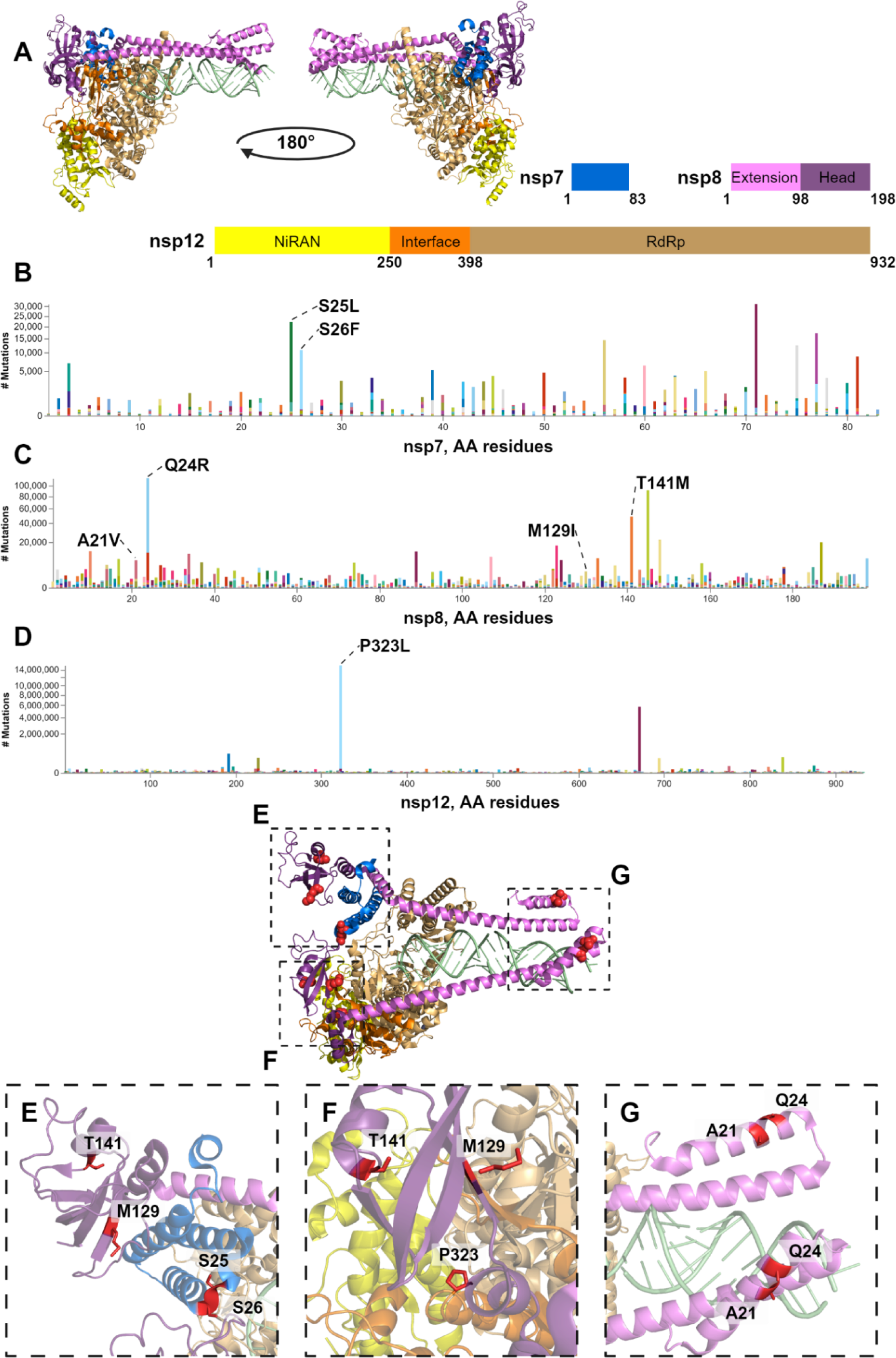
Schematic representation of SARS-CoV-2 RdRp and identified amino acid substitutions within the RdRp protein subunits. (A) A model structure (PDB id: 6yyt^12^) and domain description of the protein comprising the SARS-CoV-2 minimal RdRp complex: A single nsp7 (blue) and two nsp8 subunits are bound to nsp12. Nsp8 consists of two subdomains – a head domain (magenta) and a rod-like helical extension (pink) that stabilizes the nascent RNA duplex (green). The multidomain nsp12 is composed of the polymerase (wheat), interface (orange) and NiRAN (yellow) domains. Identified amino acid substitutions and their positions within the amino acid sequences are shown in panels B, C and D: (B) amino acid sequence of nsp7 with visualized exchanges, (C) amino acid substitutions within the nsp8 sequences, (D) identified amino acid substitutions in nsp12. The presented data were retrieved from [covidcg.org]. The whole analyzed dataset consisted of 15,243,203 sequences. Selected mutations examined in this study are labeled. The localization of amino acid residues selected for this study are visualized within the RdRp complex as red spheres with close-up views at panels E, F and G: (E) view of the nsp7 and nsp8b head domain, (F), selected amino acid residues in nsp8a head domain and nsp12 interface domain, (G) close up view of the nsp8 extension domains.

Mutations are the driving force behind the evolution and adaptation of RNA viruses. The evolution of mutant variants is closely linked to the pathogenesis and diversity in the viral population^15^. SARS-CoV-2 thus represents a perfect example of a rapidly evolving RNA virus. Five variants of concern (VOCs), designated Alpha, Beta, Gamma, Delta and Omicron, have been described since the first appearance of SARS-CoV-2 in December 2019. These variants were primarily classified based on mutations in the spike protein (S). Due to its role in immune response, target recognition, and cellular entry, the mutations in S proteins are directly involved in virulence and the infectious cycle. However, despite occurring regularly, mutations in nonstructural and accessory proteins have been studied to a much lower extent^16–18^.

Previous research on SARS-CoV-2 RdRp mutations has been focused mainly on nsp12^19, 20^, although Subissi et al, 2014^11^ demonstrated that mutations in accessory subunits, nsp7 and nsp8, can significantly influence virus fitness. Some of the polymorphisms of both nsp7 and nsp8 have been explored *in silico*, to predict their potential impact on the RdRp complex ^21, 22^. Despite the key role of RdRp in the virus life cycle, its significance in the development of drugs against COVID-19 ^23, 24^ and recognizing the potential for amino acid changes in viral enzymes to induce drug resistance ^25–27^, our knowledge of polymorphisms in the RdRp proteins remains rather limited.

To address this, we conducted a comprehensive characterization of representative naturally occurring mutations in nsp7, nsp8 and nsp12. We examined the biophysical properties of these proteins, assessed the overall enzymatic activity of the RdRp complex variants, and determined how these mutations affect the sensitivity of RdRp to Remdesivir®.

## Results

### 1. Selection of representative mutations in nsp7, nsp8 and nsp12

To select the representative mutations in RdRp subunits, we analyzed the natural amino acid substitutions occurring in nsp7, nsp8, and nsp12 over the period from January 2020 to June 30, 2022, covering all five SARS-CoV-2 VOCs (Fig 1 B, C, D). This search was based on the data in the public database [covidcg.org], encompassing a total of 15,243,203 sequences. Despite their general conservation, as indicated by data from the public database, numerous natural substitutions in RdRp accessory subunits, nsp7 and nsp8, as well as in nsp12, were observed during the pandemic. Using an experimental RdRp structure PDB ID:6yyt^12^ (Fig 1, A), we focused on amino acid residues at the interfaces that are involved in protein-protein interactions within the RdRp complex. We also considered the possibility of the mutations significantly altering the chemical properties of the respective residue.

We observed that the most frequent mutations in the nsp7 sequence were primarily located in the N-terminal region spanning residues 24-26, and in the C-terminal region, specifically involving residues 56, 71, and 77. (Fig 1, B). Due to the limited structural characterization of this nsp7 region^12, 29–31^, we focused on the residues close to the loop connecting helices 1 and 2: S24, S25, and S26 (Fig 1, E) and situated at the top of the RdRp complex near nsp12 and nsp8a. Two relatively frequent amino acid substitutions – S25L and S26F – have been identified in this region. Both of these replace the serine residue with a hydrophobic one, potentially influencing the interaction dynamic of the loop.

In nsp8, Q24 is the most frequently altered amino acid residue. It is located in the short *α*-helix at the end of the extension domain (Fig 1, C, G). Interestingly, in existing SARS-CoV-2 variants, this amino acid residue is most commonly replaced by positively charged amino acids, especially arginine. Apart from this Q24R substitution, we also selected a more subtle, yet frequent substitution, A21V (Fig 1, C, G). The head domain of nsp8 exhibits several threonine mutations, including T141, T145, or T148 (Fig 1, C). Among these, we particularly focused on T141M, which is the second most prevalent mutation within this subdomain with its position in close proximity to nsp12 (Fig 1, F). Another area of interest was the more conserved region of the nsp8 head subdomain where the M129I substitution occurs. This substitution introduces a highly hydrophobic residue at the nsp8a:nsp12 interface (Fig 1, F). Intriguingly, the M129I substitution in nsp8 was exclusively identified in sequence variants lacking the currently prevailing P323L substitution in nsp12, as reported by June 30, 2022, comprising 99.18 % of all analyzed nsp12 sequences (Fig 1, D).

The P323 residue is located directly at the interface between nsp12 and nsp8a (Fig 1, F), making it a critical site for examining interactions within the RdRp complex. The impact of this substitution on RNA polymerase activity and RdRp complex formation has been previously studied^20^. We included the P323L substitution not only to assess its direct impact on the formation and stability of the RdRp complex but also to evaluate whether mutations in accessory subunits nsp7 and nsp8 can influence the RdRp to a similar extent as mutations in the main enzymatic unit nsp12.

### 2. Impact of selected mutations on protein and RdRp complex stability

To assess the impact of the selected amino acid substitutions on the stability and polymerase activity of the RdRp, we introduced these mutations into the respective expression vectors. Subsequently, we produced and purified all proteins of interest. The wild-type (wt) nsp7 and nsp8 along with their mutant variants; nsp7 S25L and S26F and nsp8 A21V, Q24R, M129I, and T141M, were produced in *E. coli*.

Although the bacterial expression of nsp12 has been established for various purposes^32, 33^, it was also reported to potentially yield inactive or misfolded proteins, insufficient for functional studies^34^. Therefore, we produced both the wt and P323L variants of nsp12 in the *sf9* insect cell line. The mutagenesis of nsp8 and nsp12 did not impact the established purification protocol for wt proteins, nor did it significantly affect the yield. However, the purification of nsp7 variants with introduced hydrophobic residues required optimization especially due to their decreased solubility. We enhanced solubility for S25L and S26F variants by adding a mild detergent to the lysis buffer, though the yield was lower in comparison to the wt (Fig 2, A). Moreover, both mutant variants of nsp7 precipitated at higher concentrations.

**Figure 2.**
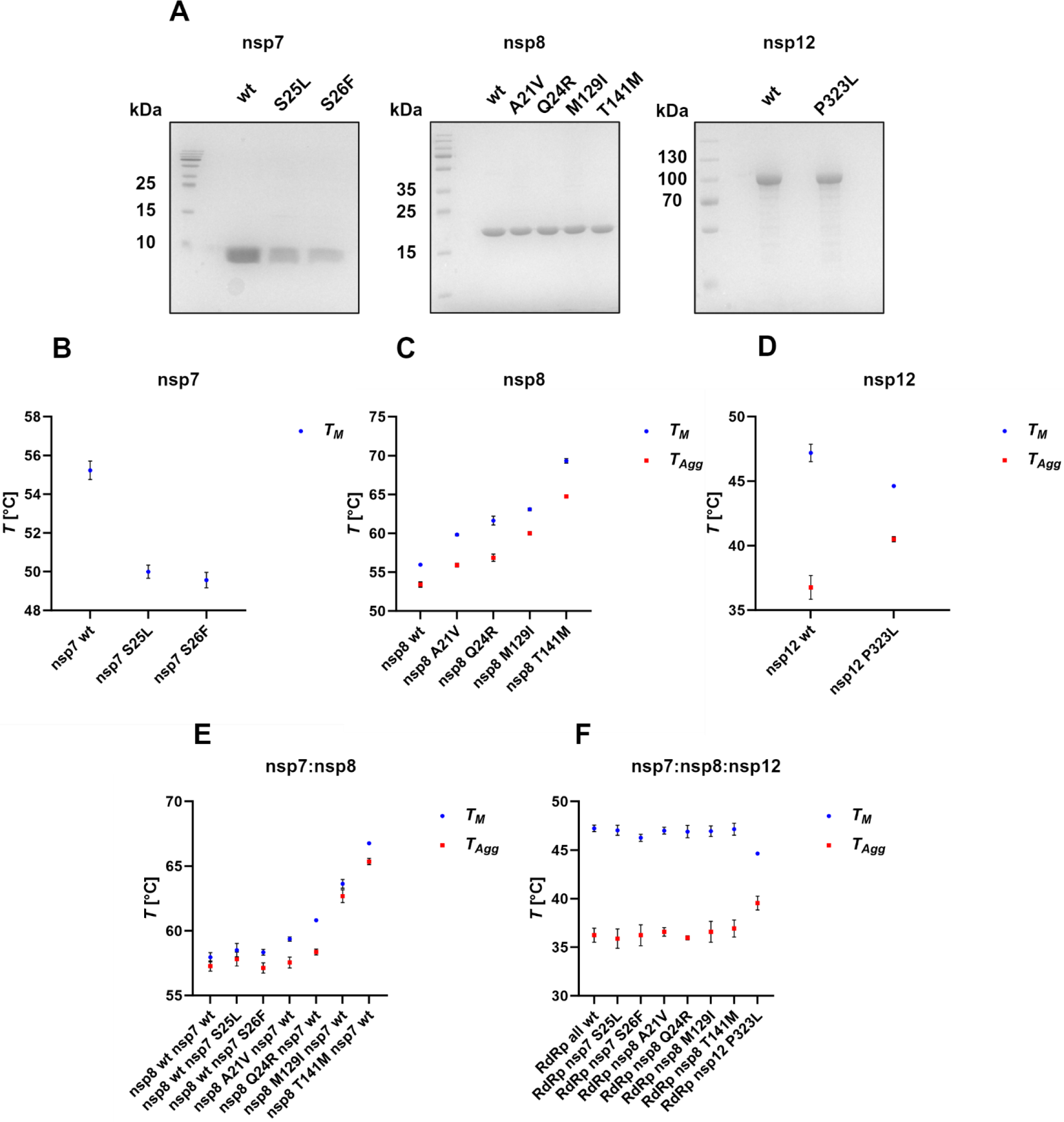
Assessment of protein stability of individual RdRp proteins and their complexes. (A) SDS-PAGE analysis of purified nsp7, nsp8, and nsp12 variants. The thermal stability (TS) of individual RdRp proteins is shown in panels A, B, C: (A) TS of S25L and S26F of nsp7 mutants, (B) TS of nsp8 mutants (C), TS of nsp12 wt and P323L. (D) TS of heterodimeric nsp7-nsp8 complexes formed by wt and indicated nsp7 and nsp8 mutants was assessed. (E) TS of nsp12 mixtures with nsp7 and nsp8 variants in a 1:1:2 ratio (nsp12:nsp7:nsp8). The tested complexes were formed by mixing two wt components with one mutant component, e.g., the RdRp nsp7 S25L complex was formed by wt nsp12, wt nsp8, and S25L variant of nsp7. The melting temperature (*T_M_*), shown in blue, determined using nanoDSF is represented by the inflection point in the 330/350 nm ratio curve. The turbidimetrically assessed aggregation temperature (*T_Agg_*), shown in red, represents the initial temperature at which the aggregation starts.

Subsequently, the stability of individual proteins and their complexes was assessed. Melting temperatures (*T_M_*) and the temperatures of the aggregation onset (*T_agg_*) were determined using a thermal unfolding assay employing nano-differential scanning fluorimetry (nanoDSF) and back-reflection turbidimetric measurement, respectively, for each mutant. In nsp7 protein, the substitution of serine at positions 25 and 26 with either leucine or phenylalanine led to its destabilization, as evidenced by a decrease of *T_M_* by approximately 4.5 °C for both S25L and S26F compared to wt (Fig 2, B, blue dots). No increase in turbidity was observed during the measurements, suggesting that none of the nsp7 variants aggregate as a result of thermal denaturation. In contrast, all analyzed amino acid substitutions in nsp8 stabilized the protein (Fig 2, C blue dots). Moreover, *T_agg_* (Fig 2, C red dots) was similarly affected as *T_M_*, confirming a direct shift in the thermal denaturation profile. The substitutions in the rod-like extension domain of nsp8, i.e. A21V and Q24R, increased the *T_M_* by 3.9 °C and 5.7 °C, respectively. M129I and T141M, located within the head domain, enhanced the stability to a greater extent. Notably, T141M had the most significant effect, increasing the *T_M_* by 13.4 °C. Concerning the nsp12 variants, the melting temperature of nsp12 P323L mutant was lower compared to the nsp12 wt; however, its aggregation temperature was proportionally higher (Fig 2, D). This indicates that the P323L variant of nsp12 undergoes a faster denaturation that starts at a higher temperature. For a larger multidomain protein like nsp12, such a shift in thermal denaturation profile typically indicates more specific denaturation. Based on these data, we concluded that the P323L substitution had a stabilizing effect.

It has been shown that nsp7 and nsp8 also form a stable heterodimeric complex, which can further assemble into larger structures, such as tetramers^31, 35^, or even hexadecamers, as shown for SARS-CoV^36^. We prepared combinations of nsp7 and nsp8 wt and mutant protein complexes in a 1:1 molar ratio, where each combination included one wt and one mutant protein, and assessed their thermal stability (Fig 2, E). No significant shift of melting temperature was observed for the complexes of wt nsp8 with destabilizing nsp7 variants whereas for nsp8 mutant variants with wt nsp7, the thermal shift was similar to individual nsp8 subunits (Fig 2, C, E).

To determine the effect of selected mutations on the stability of minimal RdRp complex, we combined wt nsp12 with each nsp7 individually, and then also with each variant of nsp8 mutant variants, as well as P323L nsp12 with wt nsp7 and wt nsp8 in a 1:1:2 ratio (Fig 2, F). No significant shifts in *T_M_* and *T_agg_* were observed, that could be attributed to substitutions in either nsp7 or nsp8. Interestingly, the same stabilizing effect observed in individual nsp12 P323L was also evident when it was combined with wt nsp7, and wt nsp8 in mixtures (Fig 2, D, F).

### 3. Effect of mutations on protein-protein interactions and RdRp complex formation

The results of thermal stability experiments indicated that the selected amino acid substitutions do not significantly disrupt protein-protein interactions within the RdRp complex. Nonetheless, proper complex formation is extremely important as nsp12 gains most of its activity only upon binding to nsp7 and nsp8^37^. Therefore, we analyzed the impact of substitutions on RdRp complex formation in more detail.

Firstly, we tested the ability of the prepared wt proteins (Fig 2, A) to form a complete dsRNA-RdRp complex using an analytical size exclusion chromatography (SEC) as described by Hillen et al., 2020^12^. The minimal RdRp complex was formed by combining wt nsp7, nsp8, and nsp12 in a 1:2:2 molar ratio, along with a short double-stranded RNA with a 10 nucleotide (nt) overhang (Fig 3, A) in a 1:1 molar ratio (dsRNA:nsp12). The proteins were detected spectrophotometrically at 280 nm and RNA at 260 nm (Fig 3, B). In addition, the peak fraction was analyzed by SDS-PAGE (Fig 3B, inlet) confirming the presence of a complex comprising all three components.

**Figure 3.**
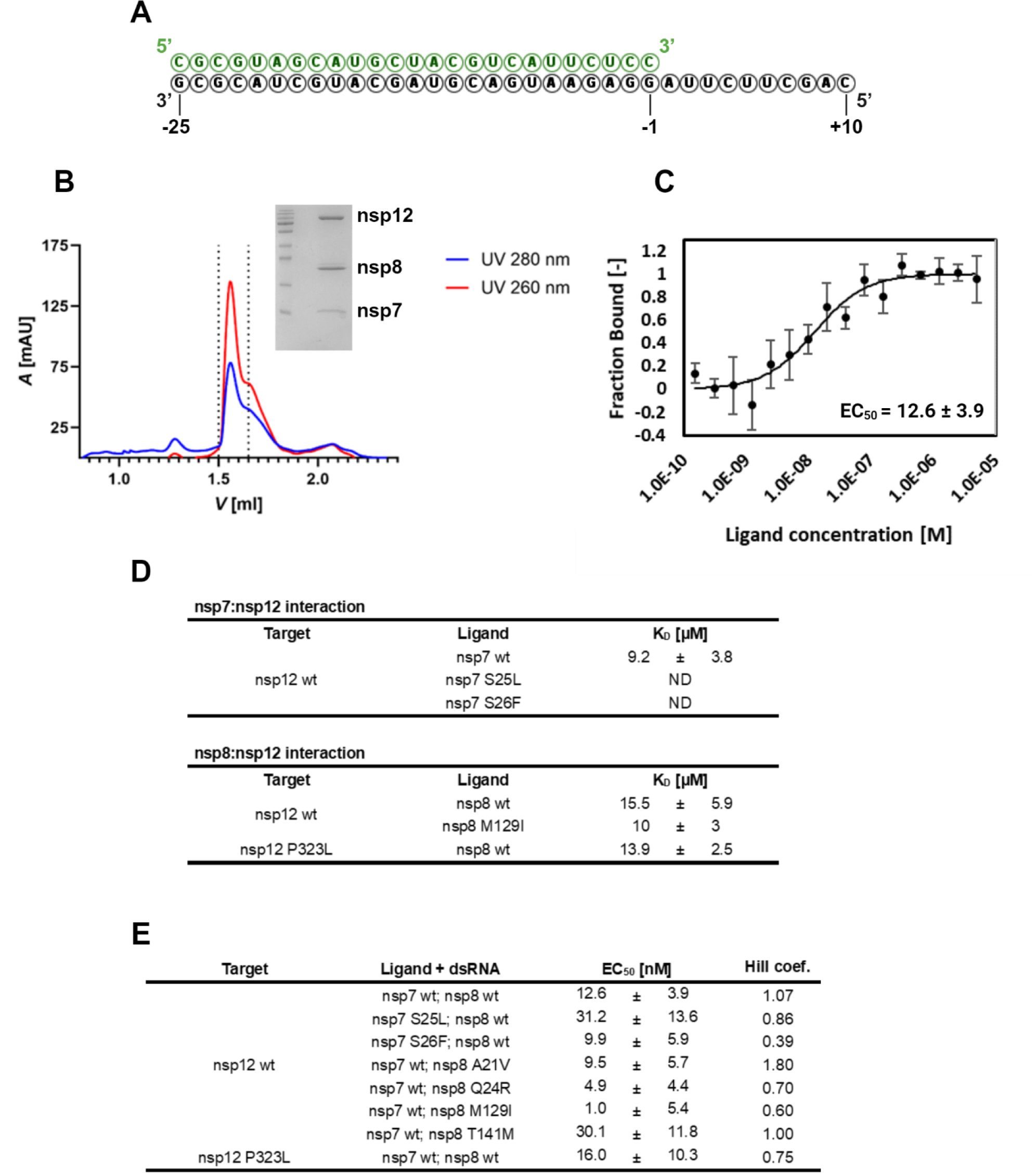
Verification of wild-type RdRp assembly with dsRNA, protein-protein interactions and RdRp assembly parameters. (A) dsRNA used for the analysis of dsRNA-RdRp complex formation. (B) Analytical size exclusion chromatography of a preformed dsRNA-RdRp complex. The complex was formed by combining individual wt components of the minimal RdRp complex, nsp7, nsp8, and nsp12 in a 1:2:2 molar ratio with the dsRNA (A) in a molar ratio of 1:1 (nsp12:dsRNA). Proteins were detected at 280 nm (blue) and RNA at 260 nm (red). The peak fraction was additionally evaluated by SDS-PAGE (inlet). (C) MST analysis of the dsRNA-RdRp complex. A subcomplex consisting of wt nsp7, wt nsp8 and dsRNA (A) in a 1:2:1 molar ratio (nsp7:nsp8:dsRNA) was used as a ligand for the wt nsp12 target. The Hill model for dose-response curve fitting was utilized to determine EC_50_. (D) Determined dissociation constants (K_D_) for nsp7:nsp12 and nsp8:nsp12 interactions by MST. (E) Experimental parameters of the assembly of dsRNA-RdRp complexes with different subunit variants determined by MST with Hill model for dose-response curve fitting.

By using a microscale thermophoresis (MST) assay, we further investigated and quantified the impact of amino acid substitutions in the RdRp components on their mutual interactions and minimal dsRNA-RdRp complex formation. Initially, we analyzed the binding affinity of nsp12 to both nsp7 and nsp8 (Fig. 3, D). The dissociation constant (K_D_) for wt nsp12 and wt nsp7 was about 9µM. However, the K_D_ for the S25L and S26F could not be determined under the same experimental conditions due to insufficient working concentration of nsp7 caused by precipitation. The K_D_ reflecting the interaction between nsp12 and nsp8 was determined to be in micromolar range both for wt and mutants with substitutions at nsp12 and nsp8 interface, specifically, nsp8 M129I and nsp12 P323L. Notably, neither of the two analyzed substitutions significantly affected the interaction (Fig 3, D).

To evaluate the impact of amino acid substitutions on the assembly and stability of the whole minimal RdRp complex with dsRNA, we used a protocol published by Kim et al., 2023^20^. The subcomplex assembled by combining nsp7, nsp8, and RNA in a molar ration 1:2:1, was used as a ligand with nsp12 serving as a target molecule in the MST assay (Fig 3, C). In these MST experiments, we used the same dsRNA as employed in the aforementioned SEC analysis for the complex formation (Fig 3, A). Instead of K_D_ mode dose-response curve fitting, we employed the Hill model for multivalent interactions, which provides information about the cooperativity of the interaction and the half-maximal effective concentration (EC_50_), thus representing the concentration at which 50 % of the labeled nsp12 binds to ligands^38^. The EC_50_ of the RdRp wt complex was in the nanomolar range (Fig 3, C), in contrast to the micromolar range K_D_ observed for nsp7-nsp12 or nsp8-nsp12 interactions (Fig 3, D). We next investigated the capability of nsp7, nsp8 or nsp12 variants to form the minimal RdRp complex and quantified the interaction parameters for each mutant. The alterations in complex formation characterized by EC_50_ were predominantly negligible (Fig 3, E). However, the increased EC_50_ observed for the S25L substitution in nsp7 and T141M in nsp8, suggests their negative effect on complex formation. Moreover, except for nsp8 A21V, all mutant variants exhibited decreased Hill coefficient, indicating negatively affected cooperativity during the assembly.

### 4. Mutations influence the overall RdRp activity

As the protein stability of all three RdRp subunits and complex formation were influenced by the amino acid substitutions, we hypothesized that the overall enzymatic activity of the RdRp complex might be affected as well. To test this hypothesis, we adapted a fluorometric RNA extension assay used for the analysis of ZIKA virus polymerase activity^39^, with adjustments for the SARS-CoV-2 RdRp^40^. Briefly, the activity assay utilizes a 20 nt RNA primer RNA annealed to a 79 nt template RNA, which includes a complementary region and a 59 nt polyU 5’ overhang (Fig 4, A). Elongation of the primer RNA was measured using SYBR® Green I, an intercalating fluorescent dye. Fluorescence was measured in real-time generating a fluorescence curve for each reaction. To compare the enzymatic activities of the protein variants, the area under the curve (AUC) was calculated for each fluorescence curve, with representative examples shown in Fig 4, B. The AUC of a reaction comprising wt nsp7, nsp8 and nsp12 served as a reference standard, set to 100%, and the results for the mutants are presented as a percentage of the wt RdRp activity. For each substitution, at least three independent measurements were conducted.

**Figure 4.**
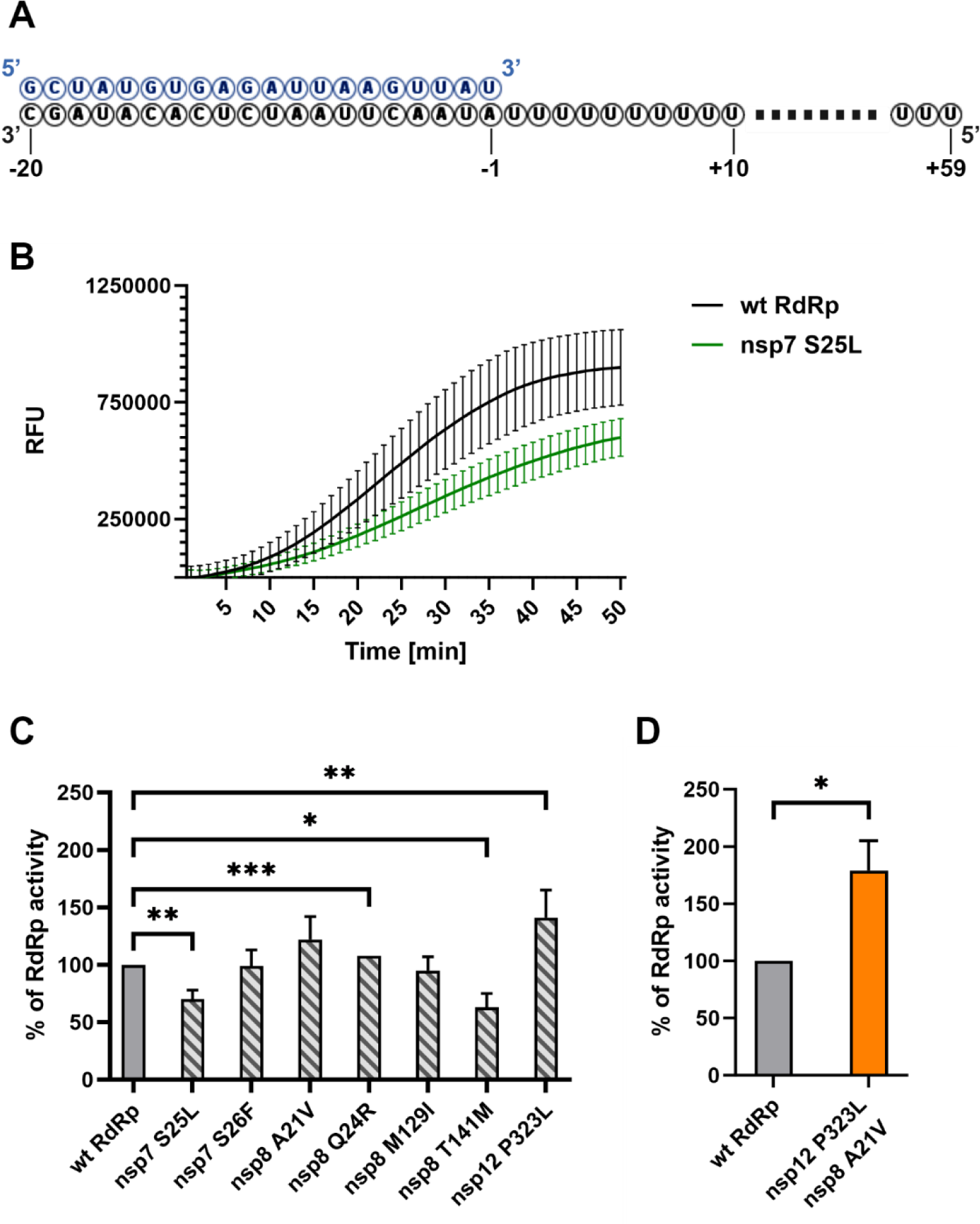
Amino acid substitutions of individual RdRp proteins alter the polymerase activity. (A) A Primed polyU RNA template used to analyze the RdRp activity. (B) Comparison of the fluorescence curves for RdRp complex of wt nsp7, wt nsp8 and wt nsp12 (labeled “wt RdRp”) and nsp7 S25L, wt nsp8 and wt nsp12 (labeled “nsp7 S25L”). (C) RNA polymerase activity of different RdRp variants compared to a combination of wt nsp7, wt nsp8 and nsp12. Each variant comprised one subunit with an amino acid substitution, e.g. nsp7 S25L represents a combination of nsp7 S25L, wt nsp8 and wt nsp12. (D) The most significant increase in RdRp activity was observed for a complex of wt nsp7, nsp8 A21V and nsp12 P323L. Results represent statistically processed data from at least three independent measurements. Outliers were excluded using a Q-test at a 90 % significance level. A paired two-tailed t-test was utilized to compare results with wt RdRp with indicated p values (* = p ≤ 0.05; ** = p ≤ 0.01; *** = p ≤ 0.001). The % of RdRp activity was calculated by comparing the area under the curve (AUC) with the AUC of wt RdRp. RFU – relative fluorescence units.

RdRp activity was initially measured using nsp7-nsp8-nsp12 complexes, each containing a single protein with an amino acid substitution. The S25L substitution in nsp7 negatively affected complex formation, leading to a decrease in RdRp activity to 70.2 ± 8.3 %. In contrast, the S26F substitution in nsp7 did not significantly affect the enzymatic activity. The impact of substitutions in nsp8 on RdRp activity varied depending on the location of the respective mutated residue. Substitutions within rod-like extensions of nsp8 slightly activated the RdRp, specifically, Q24R increased the activity by 7.8 ± 0.3 %. Conversely, the M129I substitution in the head domain of nsp8 had a negligible effect on RNA polymerase activity, and the exchange of T141 per methionine decreased the activity to 63.1 ± 12.1 % compared to the wt RdRp (Fig 4, C).

The P323L variant of nsp12 was previously proposed to enhance the RNA polymerase activity^20^. Using the fluorometric method, we also observed an increase in the activity for the P323L nsp12 variant to 141.3 ± 24.0 % relative to the wt nsp12 (Fig 4, C). This particular nsp12 variant was first identified in February 2020^41^ and during the pandemic, was found to co-occur with nearly every substitution in nsp7 and nsp8, except for M129I in nsp8 [covidcg.org]. Therefore, we also evaluated the relative activity of RdRp complexes consisting of the nsp12 P323L variant in combination with various nsp7 and nsp8 variants. A notable synergistic effect was observed especially between nsp12 P323L and nsp8 A21V mutants (Fig 4, D), which elevated the overall RdRp activity to 178.9 ± 25.7 % compared to the wt RdRp complex.

### 5. Mutations alter the response to Remdesivir®-triphosphate

Interfering with RNA replication and RdRp targeting is a common strategy in antiviral drug development^42^. Remdesivir®, initially developed as a treatment for the Ebola virus^43^, has demonstrated inhibitory activity against multiple RNA viruses^44^, including SARS-CoV-2^45, 46^. Moreover, it is also one of the drugs approved for use against COVID-19^47^. Remdesivir® functions as a prodrug that is biotransformed after administration and converted into an adenosine-triphosphate analog (Fig 5, A). Its mode of action involves incorporation into the nascent RNA strand instead of adenosine, leading to the premature termination of RNA synthesis. The termination occurs due to steric hindrance after adding three subsequent nucleotides, translocating Remdesivir®-monophosphate (RMP) to the –4 position^45^. As the investigated mutations induced significant changes in RdRp activity, we sought to determine whether amino acid substitutions of RdRp components could affect the inhibitory efficacy of Remdesivir®. As a positive control for these experiments, we first assessed the inhibition of wt RdRp, by monitoring the elongation of the dsRNA (Fig 4, A) in the presence of 100µM Remdesivir®-triphosphate (RTP) (Fig 5, B). The residual activity (RA) was determined by comparing the AUC of the reaction in the presence of RTP to the AUC of the uninhibited reaction.

**Figure 5.**
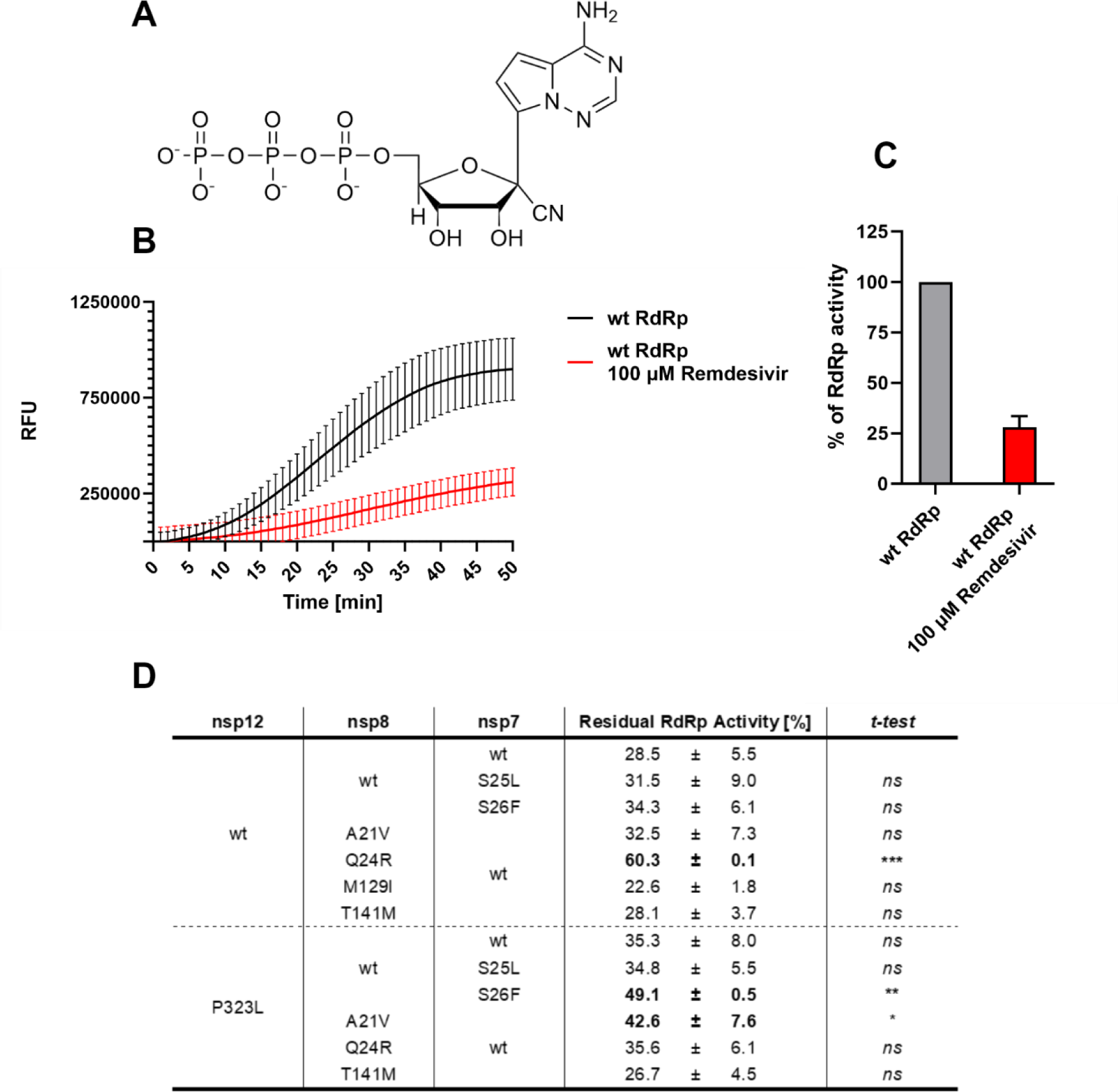
Fluorometric assessment of inhibitory activity of Remdesivir®-triphosphate. (A) Structure of Remdesivir®-triphosphate (RTP). (B) Comparison of fluorescence curves recorded for wt RdRp (black) and for RdRp in the presence of 100µM RTP (red). (C) Calculated residual activity (RA) of wt RdRp in % of RdRp activity for reaction with 100µM RTP (red) compared to uninhibited reaction (gray). (D) Residual RdRp activities in % measured for various nsp7:nsp8:nsp12 combinations in the presence of 100µM RTP. Results represent statistically processed data from at least three independent measurements. Outliers were excluded using a Q-test at a 90 % significance level. The % of RdRp activity was calculated by comparing the area under the curve (AUC) with the AUC of wt RdRp. A paired two-tailed t-test was utilized to compare RAs of various RdRp variants with RA of RdRp composed of wt nsp7, wt nsp8 and nsp12 with indicated p values (*ns =* p > 0.05; * = p ≤ 0.05; ** = p ≤ 0.01; *** = p ≤ 0.001). RFU – relative fluorescence units.

The respective RAs were subsequently measured for RdRp complexes, encompassing all nsp7 and nsp8 variants combined with either wt or the P323L nsp12 variant. The RA of the wt RdRp, which was found to be 28.5 ± 5.5 % (Fig 5, C), served as a reference for statistical comparison with each of the tested combinations (Fig 5, D). Notably, three combinations exhibited a significant increase in RA, two of which comprised the nsp12 P323L. Specifically, the combination of nsp12 P323L with nsp8 A21V showed an RA 42.6 ± 7.6 %. Another combination involving S26F substitution in nsp7, resulted in an RA of 49.1 ± 0.5 %. Interestingly, the most Remdesivir®-resistant RdRp variant was a combination of wt nsp12 with nsp8 Q24R, exhibiting nearly double RA compared to that of the wt RdRp. None of the RdRp complexes exhibited statistically significant increase in sensitivity to RTP. The lowest RA (22.6 ± 1.8 %) was observed for the RdRp complex comprising wt nsp7, nsp8 M129I and wt nsp12.

## Discussion

Following the emergence of COVID-19, the extensive transmission of SARS-CoV-2 led to the development of numerous variants carrying mutations which resulted in amino acid substitutions also identified in the proteins constituting the SARS-CoV-2 RdRp complex (Fig 1) potentially influencing virulence or resistance. In this study, we selected representative mutations in RdRp subunits, specifically S25L and S26F substitutions in nsp7; A21V, Q24R, M129I and T141M in nsp8 and a dominant substitution P323L in nsp12. Corresponding proteins were cloned, expressed and purified for a multiscale characterization of the phenotypic impact of the mutations.

The substitutions within all three subunits of the SARS-CoV-2 RdRp complex significantly influenced their thermal stability. The scale of thermal shift observed in both nsp7 and nsp8 mutants (Fig 2 B, C) suggests that the effect of the substitutions is not solely intrinsic, but most likely due to the changes in the interactions ultimately leading to the formation of the RdRp complex. Unfortunately, reports on nsp7 self-interactions have been diverse^30, 48^, making it challenging to determine whether the effect is due to a reduced multimerization. On the contrary, nsp8 was shown to form homodimers and homotetramers in solution^30, 49^, indicating that the thermal shift could be induced as a result of enhanced multimerization. Regarding the nsp7:nsp8 complexes, thermal stability was primarily affected by the nsp8 subunit variant (Fig 2, C, E). This suggests that the effect of various substitutions might extend from nsp8 homomultimers to nsp7:nsp8 heteromultimers. Moreover, none of the analyzed amino acid substitutions is located near the nsp7:nsp8 interface^35^, implying that these substitutions should not interfere with the complex formation as observed for various nsp7 variants (Fig 2, E). While the complex of nsp7 with the C-terminal domain of nsp8 is structurally well characterized^36^, much less is known about the position of the N-terminal extension within these complexes. Additionally, the structure of nsp8 homomultimers has not been determined yet. It is known that the N-terminal extension may adopt multiple conformations, as shown for the SARS-CoV nsp7:nsp8 hexadecamer^50^ or the Feline coronavirus nsp7:nsp8 heterotrimer^30^. In addition, Wilamowski et al. 2021^30^, proposed possible interactions between N-and C-termini of nsp8 in solution, combining SAXS and SANS techniques. These interactions might be enhanced by selected substitutions in nsp8, potentially increasing the stability of both nsp8 homomultimers and nsp7:nsp8 complexes.

As indicated by the thermal unfolding assay, selected amino acid substitutions did not affect the RdRp complex formation (Fig 2, F). To verify these findings, we quantified the protein-protein interactions using an MST assay. The interactions between nsp7:nsp12 and nsp8:nsp12 showed a K_D_ in micromolar range. In contrast, the formation of the RdRp complex with both nsp7 and nsp8 subunits along with dsRNA occurred within the nanomolar EC_50_. This suggests that the presence of all RdRp subunits together with the dsRNA greatly enhances the individual interactions that lead to the formation of the dsRNA-RdRp complex, which is consistent with previously published data by Kim et al., 2023^20^. However, our results slightly differed from that of Kim et al., 2023. Firstly, the EC_50_ in our study indicated greater affinity during the assembly of wt RdRp and secondly, we found no significant difference in complex formation between wt and P323L nsp12 variants. These discrepancies could be attributed to the differences in the nsp12 purification protocol. The protocol used in this study consisted of multiple steps including tag cleavage, while Kim et al., 2023 utilized nsp12 purified with a single step NiNTA chromatography.

Our results showed that even a single amino acid substitution in any of the protein subunits of SARS-CoV-2 RdRp could alter the enzymatic activity of the complex (Fig 4, C). Noteworthy, substitutions within nsp7 and nsp8 did not change the RdRp activity as substantially as P323L in nsp12, and even in comparison with other studied mutations in nsp12^19^, the impact was less significant. As anticipated, the differences in enzymatic activity align with alterations in the biophysical properties of the proteins. For instance, S25L in nsp7 and T141M in nsp8 showed higher EC_50_ values in MST experiments, subsequently exhibiting a notable decrease in RdRp activity. Conversely, nsp7 S26F and nsp8 M129I, which largely retained the relative RdRp activity, exhibited comparable EC_50_ values to their respective wt variants. Intriguingly, both of these mutations decreased the Hill coefficient of the complex assembly, suggesting that the cooperativity of the assembly is unrelated to the overall enzymatic activity. Concerning the mutations in the N-terminal rod-like extension domain of nsp8, both A21V and Q24R mutants tended to increase the catalytic activity of the RdRp; however, with varying effects on assembly parameters. This increase in activity does not seem to depend on the enhanced ability of complex formation. Instead, it is plausible that these substitutions promote intermolecular interactions within the helix-turn-helix motif at the N-terminus (Fig 1, G)^12, 51^, as suggested by a slight increase in thermal stability observed for these nsp8 variants (Fig 2, C, E). This finding is similar to the increase in activity observed by Subissi et al., 2014^11^ for SARS-CoV nsp8 mutant with S11A substitution located within this motif. Moreover, some of our experimental results differ from the *in silico* analyses of amino acid substitutions in subunits of SARS-CoV-2 RdRp. For example, Reshamwala et al., 2021^22^ predicted that M129I in nsp8 and S26F in nsp7 might potentially benefit the RdRp complex by increasing the surface complementarity of the subunits. However, we observed that S26F and M129I had no significant effect on either complex formation or catalytic activity. In contrast, T141M substitution in nsp8 was predicted to destabilize the RdRp complex by Senthilazhagan et al., 2023^21^, which was confirmed by our data. Both mentioned studies suggested that substituting S25 in nsp7 per leucine could stabilize the RdRp, but our results showed the opposite (Fig 3, E, Fig 4, C). Finally, while the P323L substitution in nsp12 is regarded as a stabilizing mutation, our data revealed that although it stabilized the protein itself (Fig 2, D, E), it did not enhance the protein-protein interactions (Fig 3, D, E).

Our experimental results support the overall favorable nature of the P323L substitution in nsp12 as previous research indicated that SARS-CoV-2 variants carrying this mutation in RdRp exhibit higher infectivity and enhanced selective advantage^20, 52, 53^. Additionally, the nsp12 P323L mutation is also linked to an increased mutation rate during RNA replication, potentially accelerating virus evolution^41^. Compared to nsp8 M129I mutation, which according to sequence analysis [covidcg.org] is strictly associated with wt nsp12, representing thus an alternate evolutionary pathway for the virus, the substitution of leucine for P323 in nsp12 proved to be more advantageous. Mutations in nsp7 and nsp8 were generally identified in a wide range of isolates without any apparent mutual relation or phenotypic effect, suggesting their evolution might be presumably random. Notably, among these mutations, only the nsp7 S25L variant, which negatively impacted RdRp function, has been linked to a specific locally predominant SARS-CoV-2 lineage – the B.1.497 subvariant – circulating in South Korea in 2020^54, 55^. Furthermore, considering the observed impacts on the thermal stability of individual RdRp subunits, it is also noteworthy that there might be potential implications for the course of COVID-19. According to Kim et al. 2023^20^, the nsp12 P323L mutation affects the RdRp activity at different temperatures, directing the virus to replicate more rapidly in the upper respiratory tract. Similar effects might thus result from the analyzed mutations in this study.

As the SARS-CoV-2 RdRp is a crucial therapeutic target, we tested how various substitutions impact the inhibitory effect of the FDA-approved drug Remdesivir® functioning as an RNA chain terminator. Previous reports have highlighted the superior selectivity of RTP compared to adenosine triphosphate^46^, attributed to a cyano moiety (Fig 5, A) fitting into a hydrophobic pocket in nsp12, defined by residues T687, N691 and S759^32^. Furthermore, the replication stalling by RMP results from a steric clash with S861 after RMP translocation to –4 position^45^. Both RTP binding and stalling thus occur within the polymerase domain suggesting that the substitutions we analyzed should not directly interfere with the RTP inhibition mechanism. However, our study identified that multiple RdRp variants, namely combinations of wt:Q24R:wt, wt:A21V:P323L and S26F:wt:P323L (nsp7:nsp8:nsp12), exhibit a reduced rate of relative inhibition (Fig 5, D). The RMP-induced stalling results in blockage of translocation, prompting nsp12 to adopt the pre-translocated conformation^45, 56^; however, this state was reported to not be definitive as the stalling can be overcome at higher NTP concentrations^40^. The regulation of RdRp translocation was additionally linked with the nsp7:nsp8 interaction, with the potential involvement of the N-terminal extension of nsp8^57^. Considering that the P323L was not sufficient to reduce the RTP inhibition, we hypothesize that the observed effect could result from differences in translocation regulation caused by substitutions in nsp7 and nsp8. However, a more detailed understanding of how specific amino acid exchanges lead to nucleotide analog resistance requires further investigation. Moreover, these mutations may also represent an obstacle in the designing of non-nucleoside or allosteric RdRp inhibitors. Particularly, the nsp8-nsp12 interaction interface and the interface domain of nsp12 have been identified as potential hot spots for small molecule inhibitors, with the binding site in nsp12 directly involving the P323 or surrounding amino acid residues^58–60^.

In conclusion, our findings elucidate the impact of naturally occurring amino acid substitutions within the RdRp proteins of SARS-CoV-2. Understanding these changes is crucial, as they can significantly influence interaction dynamics of the RdRp complex and the enzymatic activity, consequently altering the RdRp sensitivity to drugs such as Remdesivir®. This knowledge is particularly important for the development of antiviral strategies targeting RdRp.

## Methods

### 1. Preparation of expression vectors

Expression vectors encoding 6×His-TEV-nsp7 (#154757), 6×His-TEV-nsp8 (#154758) and 6×HisMBP-TEV-nsp12 (#154759) were obtained from AddGene^12^. Efficient Mutagenesis Independent of Ligation (EMILI)^28^ was utilized to introduce mutations in the respective sequences. Briefly, primers encoding the mismatch mutations (Tab S1) were used in an inverse PCR to amplify the entire expression vector. Subsequently, the methylated template DNA was digested with DpnI for 90 minutes at 37 °C. The mixtures were then incubated with T4 DNA polymerase for 5 minutes at 22 °C to generate sticky ends. To inactivate T4 DNA polymerase, EDTA was added to the final concentration of 2.5mM and the temperature was increased to 72 °C for 20 minutes. The whole reaction mixture was then used for the transformation of *Escherichia coli* DH5α competent cells. Finally, selected colonies were used to amplify the vector with introduced mutations. Each expression vector was verified by Sanger sequencing.

### 2. Protein production and purification

#### a) nsp7 and nsp8

6×His-TEV-nsp7 and 6×His-TEV-nsp8 variants were separately expressed in *E. coli* BL21 DE3 CodonPlus RIL. The cells were grown in LB medium at 37 °C to OD_600_ 0.6. Subsequently, the cultures were cooled to 18 °C, induced with IPTG at a final concentration of 0.4 mM and protein expression continued for 16 hours at 18 °C. Cells were then separated from the media by centrifugation (3,000 × g, 15 minutes), resuspended in nsp7/8 Lysis buffer (50 mM Na-HEPES pH 7.4, 300 mM NaCl, 30 mM imidazole, 10 % glycerol, 2 mM β-mercaptoethanol) and disrupted by high pressure using a One Shot Cell Disruptor (Constant Systems) at 1.9 kbar. In the case of nsp7, Triton X-100 was added before the disruption to the final concentration of 0.15 %. The lysate was then cleared by centrifugation (50,000 × g, 30 minutes), and the supernatant was loaded on HisTrap™ HP (Cytiva) preequilibrated in Lysis buffer. The column was washed with High Salt buffer and Low Salt buffer containing 1000mM NaCl and 150mM NaCl, respectively. The proteins were then eluted with Elution buffer (50mM Na-HEPES pH 7.4, 150mM NaCl, 500mM imidazole, 10% glycerol, 2mM β-mercaptoethanol). Peak fractions were collected, and His-tagged TEV protease was added at a 1:10 w:w ratio. The mixture was dialyzed against 3 liters of Low salt buffer at 4 °C for 16 hours to remove imidazole. After dialysis, the sample was loaded on the HisTrap™ HP column equilibrated in Low Salt buffer to remove the uncleaved protein and His-TEV protease. Flowthrough fraction containing the cleaved protein was collected and dialyzed against 3 liters of Nsp7/8 Storage Buffer (20mM Na-HEPES pH 7.4, 150mM NaCl, 5 % glycerol, 1mM TCEP). Finally, the samples were concentrated, aliquoted, and stored at -80 °C until use.

#### b) nsp12

Bacmids containing 6×His-MBP-TEV-nsp12 variants were generated using the Bac-to-Bac™ baculovirus expression system (Invitrogen). *Sf9* cells were transfected with the bacmid DNA and baculovirus-containing cell supernatants were harvested 5 days after transfection. The resulting baculoviral stock was amplified to achieve optimal infectivity. Subsequently, a suspension of *Sf9* cells (3×10^6^ cells/ml) was infected with the baculoviral stock in a 1:100 ratio. The expression of the 6×His-MBP-TEV-Nsp12 protein was carried out for 96 hours after which the cells were harvested by centrifugation (3,000 × g, 15 minutes). To isolate the protein, the cell pellet was resuspended in Lysis buffer (50 mM Na-HEPES pH 7.4, 300mM NaCl, 30mM imidazole, 3mM MgCl_2_ 10% glycerol, 5mM β-mercaptoethanol) and homogenized utilizing One Shot Cell Disruptor (Constant Systems) operating at 1.0 kbar. Cell lysate was subsequently clarified by centrifugation at 70,000 × g for 30 minutes and ultracentrifugation at 235,000 × g for 60 minutes. The obtained supernatant was then loaded on HisTrap™ HP (Cytiva) in Lysis buffer and washed with High salt buffer containing 1,000mM NaCl. A MBPTrap™ HP (Cytiva) column was connected downstream to the HisTrap™ HP, from which the protein was then eluted with 500mM imidazole. HisTrap™ column was subsequently disconnected and the MBPTrap™ HP was washed with additional 5 column volumes of Lysis buffer. Finally, nsp12 was eluted with a buffer containing 116.9mM maltose. 6×His-TEV protease was added to the eluate in a 1:10 w:w ratio and the sample was dialyzed against 3 liters of Lysis buffer at 4 °C for 16 hours. Reverse IMAC was utilized to remove TEV protease, cleaved purification tags, and uncleaved protein. Nsp12 was further purified using heparin affinity chromatography. For this purpose, the sample was diluted to 80mM NaCl using a Dilution buffer (20mM Na-HEPES pH 7.4, 1mM MgCl_2_, 10% glycerol, 5mM β-mercaptoethanol) and loaded on HiTrap heparin HP (Cytiva) in Binding buffer (20mM Na-HEPES pH 7.4, 80mM NaCl, 1mM MgCl_2_, 10 % glycerol, 5mM β-mercaptoethanol). The protein was then eluted with a linear gradient of NaCl from 80 to 1,000 mM. The fractions containing nsp12 were pooled and dialyzed for 16 hours against 3 liters of Storage buffer (20 mM Na-HEPES pH 7.4, 300mM NaCl, 1mM MgCl_2_, 10% glycerol, 1mM TCEP). The final sample was concentrated and frozen at -80 °C as single-use aliquots.

### 3. Analytical size exclusion chromatography

RNA 25-mer 5′-CGCGUAGCAUGCUACGUCAUUCUCC-3′ and 35-mer 5′-CAGCUUCUUAGGAGAAUGACGUAGCAUGCUACGCG-3′ were first mixed in a 1:1 ratio in Annealing buffer (20mM Na-HEPES pH 8, 10mM KCl, 6mM MgCl_2_, 0.01%Triton X-100, 1mM DTT). The mixture was then heated at 95 °C for 5 minutes and stepwise cooled to room temperature to generate an RNA duplex (Assembly RNA). Wild-type nsp12 was subsequently added to the RNA in a 1:1 molar ratio whereas nsp7 and nsp8 were both supplemented in a 2:1 molar ratio. The complete mixture was incubated at 4 °C for 2 hours to allow for stable assembly. Subsequently, the RNA-RdRp complex was loaded on Superdex Increase 3.2/300 column (Cytiva) in RdRp Assembly buffer (20mM Na-HEPES pH 7.4, 100mM NaCl, 1mM MgCl_2_, 1mM TCEP). UV absorbance at 260 nm and 280 nm was monitored throughout the analysis.

### 4. Thermal unfolding assay

Nsp7, nsp8, and nsp12 variants were first diluted to a suitable concentration to facilitate the fluorescence signal – 50 µM for both nsp8 and nsp7 and 10 µM for nsp12. To analyze the nsp7:nsp8 dimers and nsp7:nsp8:nsp12 mixtures, the proteins were mixed in 1:1, or 1:2:1 ratio, respectively. Thermal unfolding parameters were then determined using a combination of nano-differential scanning fluorimetry (nanoDSF) and turbidimetry using Prometheus Panta (Nanotemper). Protein solutions were heated at 2 °C per minute from 25 °C to 95 °C. Intrinsic fluorescence (λ_ex_ = 280 nm, λ_em_ = 330 and 350 nm) and turbidity were measured throughout the process. Results were analyzed using the Panta Analysis software (Nanotemper). Melting temperature *T_M_* was collected as the first derivative of the 330/350 fluorescence ratio whereas *T_Agg_* represents the onset temperature of turbidity increase.

### 5. Microscale thermophoresis assay

Wt and P323L variants of nsp12 were labeled with NHS-fluorescent label utilizing the Protein labelling kit (Nanotemper). Labeled proteins were then diluted to either 105nM for protein-protein interactions or 52.5 nM for complex formation and utilized as a target molecule. The stock solutions of nsp7 or nsp8 variants were used as a ligand to analyze the protein-protein interactions. To evaluate complex formation, the ligand was prepared by mixing various variants of nsp7 and nsp8 with Assembly RNA in a 1:2:1 ratio (nsp7:nsp8:RNA) to the final concentrations of 10µM, 20µM and 10µM. Every nsp7:nsp8:RNA subcomplex was incubated for 15 minutes at room temperature before the next step. The respective ligands were then diluted to make 16 two-fold dilutions. For the experiment, 10 µl of each ligand dilution was mixed with 10 µl of the respective target molecule and the obtained mixtures were incubated for an additional 15 minutes and subsequently cleared by centrifugation. Each mixture was then loaded into Monolith NT.115 capillaries (Nanotemper). The measurements were conducted using the Monolith NT.115 instrument (Nanotemper) at 25°C with the setting adjusted to 20 % LED power and medium MST power. Three independent measurements were carried out for each binding experiment. The obtained data were analyzed using MO affinity (Nanotemper) using MST-on time 19 to 20 s for protein-protein interactions and 0.5 to 1.5 s for complex formation. K_D_ mode fitting was utilized to evaluate protein-protein interactions whereas complex formation was assessed through the Hill model for multivalent binding, resulting in the determination of EC_50_ and Hill coefficient values.

### 6. Fluorometric RNA extension assay

The Substrate RNA for the extension assay was prepared as follows: The RNA template 5′-U_59_AUAACUUAAUCUCACAUAGC-3′ and RNA primer 5′-GCUAUGUGAGAUUAAGUUAU-3′ were mixed in 1:1 molar ratio in Annealing buffer (20mM Na-HEPES pH 8, 10mM KCl, 6mM MgCl_2_, 0.01%Triton X-100, 1mM DTT). The RdRp activity was subsequently monitored in real-time using SYBR® Green I, which causes an increase in emitted fluorescence through binding exclusively to double-stranded nucleic acid. The reaction mixtures were prepared by mixing nsp7, nsp8 and nsp12 in a 1:3:3 molar ratio with final concentrations of 0.63µM, 1.88µM and 1.88µM along with 250µM rATP and 5µM SYBR® Green I in Reaction buffer (20mM Na-HEPES pH 8, 10mM KCl, 6mM MgCl_2_, 0.01%Triton X-100, 1mM DTT). The mixtures were then transferred to a 96-well plate and reactions were initiated by adding Substrate RNA to the final concentration 0.5 µM. The well-plate was subsequently incubated at 30 °C for 50 minutes in the thermocycler QuantStudio 5 real-time PCR system (Applied Biosystems). SYBR® Green I fluorescence was recorded each minute throughout the incubation, resulting in a fluorescence curve for each reaction. To explore the inhibition rate, Remdesivir®-triphosphate was added in the final concentration of 100µM to the reaction mixture.

### 7. RdRp activity evaluation

To evaluate the activity of the RdRp, the Area Under the Curve (AUC) was calculated as a numerical integration of each obtained fluorescence curve using the midpoint method. Therefore, the AUC represents a sum of rectangles, where the area of each rectangle is calculated by multiplying the average of two adjacent values for fluorescence signal by a step of 1 min. The calculated AUC for each variant was referred to AUC of the wild-type RdRp reaction. To analyze the inhibition rate by Remdesivir®, the calculated AUC of the inhibited reaction was referred to the AUC of the uninhibited one with the same protein variants. Data were collected from at least three independent measurements with two duplicate reactions within each experiment. Outliers were excluded by using the Q-test. A two-tailed paired T-test was utilized to statistically compare results with the respective references.

## Acknowledgment

This study was supported by the project National Institute of Virology and Bacteriology (Programme EXCELES, ID Project No. LX22NPO5103) - Funded by the European Union - Next Generation EU and by Czech Science Foundation (No. 22-17118S).

## Supplementary information

Figures S1: Example Thermal unfolding profiles

Table S1: List of oligonucleotides used for mutagenesis

## Author Contributions

MD: mutagenesis, protein production and purification, thermal unfolding assay and results evaluation, conceived ideas and experimental design, manuscript preparation

AK: RdRp activity assay, microscale thermophoresis experiments

KK: mutagenesis, protein production and purification, RdRp activity assay

AD: Fluorometric RdRp extension assay, data evaluation

AP: Thermal unfolding assay, microscale thermophoresis experiments

JP: microscale thermophoresis experiments and results evaluation

MK: RdRp activity results evaluation

TR: conceived ideas, manuscript preparation

MR: conceived ideas and experimental design, manuscript preparation All authors reviewed the manuscript.

