## Supplemental Fig. S1 and Table S1 for "Biochemical characterization of naturally occurring mutations in SARS-CoV-2 RNA-dependent RNA polymerase"

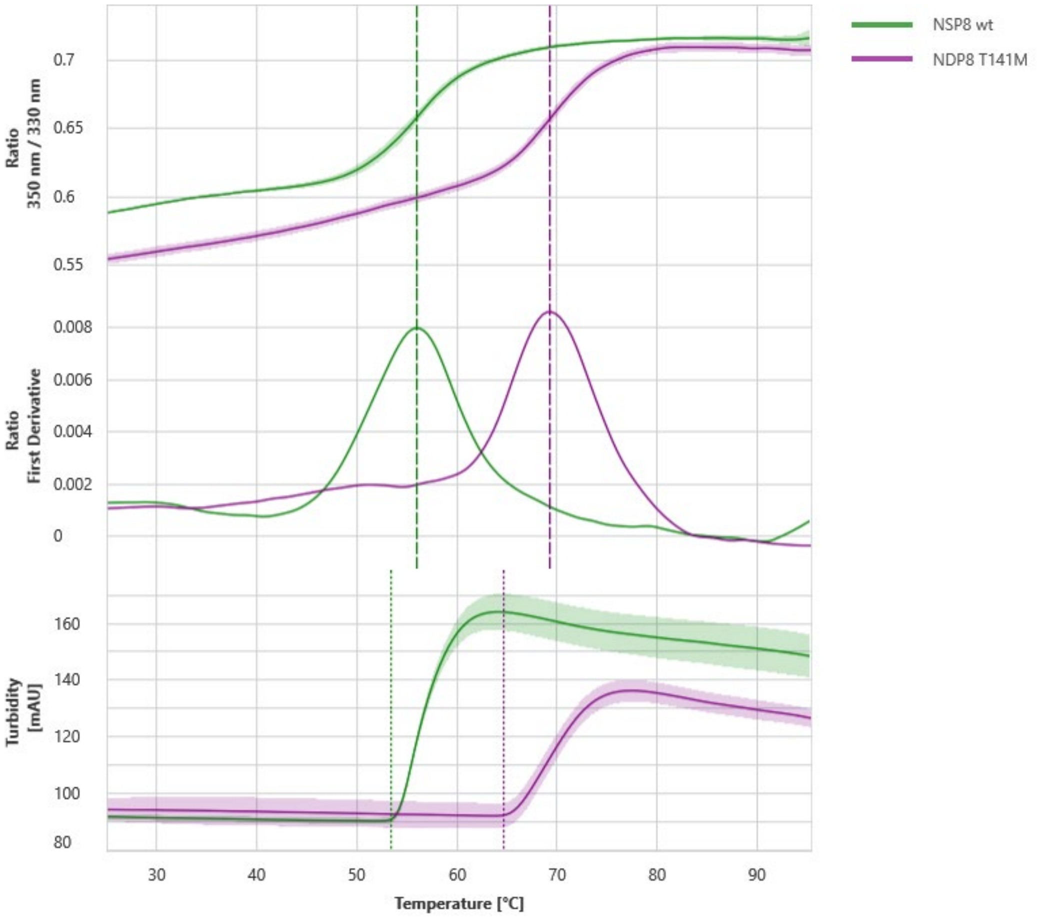


**Figure S1: Example Thermal unfolding profiles.** Thermal unfolding of nsp8 wt (green) and nsp8 T141M (violet). Ratio of fluorescence at 350 nm and 330 nm (top) and first derivative of the ratio (middle) were used to collect melting temperatures ($T_M$) indicated by dashed lines. Turbidimetric measurement (bottom) was utilized to obtain onset aggregation temperature ($T_{Agg}$) shown as dotted lines.

**Table S1: List of oligonucleotides utilized for mutagenesis**

| Oligonucleotide | Sequence |
|---|---|
| nsp12 P323L fwd | GTGTTCCCTCTAACCTCCTTCGGTCCTCTCG |
| nsp12 P323L rev | CGAAGGAGGTTAGAGGGAACACAGTGG |
| nsp7 S25L fwd | GTTGAAAGCCTCAGCAAACTGTGGGCACAGTG |
| nsp7 S25L rev | CACAGTTTGCTGAGG CTT TCAACACGCAGCTG |
| nsp7 S26F fwd | GTTGAAAGCAGCTTCAAACTGTGGGCACAGTG |
| nsp7 S26F rev | CACAGTTTGAAGCTGCTTTCAACACGCAGCTG |
| nsp8 A21V fwd | CAAGAGGTATATGAACAGGCAGTTGCC |
| nsp8 A21V rev | CCTGTTCATATACCTCTTGTGCGGTTGC |
| nsp8 Q24R fwd | CTGCCCGTTCATATGCCTCTTGTGCGG |
| nsp8 Q24R rev | GGTGCCATCACACATGTTTTTATAGGTGTTGTAATC |
| nsp8 M129I fwd | GCAAAACTGATTGTGGTTATTCCGGAT |
| nsp8 M129I rev | GAATAACCACAATCAGTTTTGCTGCGGTG |
| nsp8 T141M fwd | CAACACCTATAAAAACATGTGTGATGGCACCACC |
| nsp8 T141M rev | GGTAAAGGTGGTGCCATCACACATGTTTTTATAGG |
